# A Humanized Monoclonal Antibody against the Enzymatic Subunit of Ricin Toxin Rescues Rhesus macaques from the Lethality of Aerosolized Ricin

**DOI:** 10.1101/407817

**Authors:** Chad J. Roy, Dylan J. Ehrbar, Natasha Bohorova, Ognian Bohorov, Do Kim, Michael Pauly, Kevin Whaley, Yinghui Rong, Fernando J Torres-Velez, Ellen S Vitetta, Peter J. Didier, Lara Doyle-Meyers, Larry Zeitlin, Nicholas J. Mantis

## Abstract

Ricin toxin (RT) ranks at the top of the list of potential bioweapons of concern to civilian and military personnel alike due to its high potential for morbidity and mortality after inhalation. In non-human primates, aerosolized ricin triggers a severe acute respiratory distress characterized by perivascular and alveolar edema, neutrophilic infiltration, and severe necrotizing bronchiolitis and alveolitis. There are currently no approved countermeasures for ricin intoxication. In this report, we demonstrate the therapeutic potential of huPB10, a toxin-neutralizing humanized monoclonal antibody (MAb) against an immunodominant epitope on ricin’s enzymatic A chain (RTA). Five rhesus macaques that received intravenous huPB10 (10 mg/kg) four hours after lethal dose ricin aerosol exposure all survived the toxin challenge, as compared to control animals, which succumbed to ricin intoxication within 30 h. Antibody treatment at 12 h after ricin exposure resulted in the survival of only one of five monkeys, indicating that, in the majority of animals, ricin intoxication and local tissue damage had progressed beyond the point where huPB10 intervention was beneficial. Change in pro-inflammatory cytokine/chemokines levels in bronchial alveolar lavage fluids before and after toxin challenge successfully clustered monkeys based on survival, as well as treatment group. IL-6 was the most apparent marker of ricin intoxication. This study represents the first demonstration in nonhuman primates that the lethal effects of inhalational ricin exposure can be negated by a drug candidate and opens up a path forward for product development.

## Introduction

Ricin toxin (RT) is considered a high priority biothreat agent by the Centers for Disease Control and Prevention (CDC), the US Department of Defense (DOD), and NATO due to it’s accessibility, stability, and high toxicity especially by the aerosol route. (1, 2). In nonhuman primates (NHPs), inhalation of RT elicits the clinical equivalent of acute respiratory distress syndrome (ARDS), characterized by widespread apoptosis of alveolar macrophages, intra-alveolar edema, neutrophilic infiltration, accumulation of pro-inflammatory cytokines, and fibrinous exudate (3, 4). Ricin also damages the lung mucosa and triggers vascular leak due to direct damage to endothelial cells (5). Similar effects are observed in mice, rats and swine (6-10). RT is derived from castor beans (*Ricinus communis*) and is a 65 kDa heterodimeric glycoprotein consisting of two subunits, RTA and RTB, joined via a single disulfide bond. RTB binds to glycoproteins and glycolipids on mammalian cells and facilitates the retrograde transport of RT to the endoplasmic reticulum (ER). In the ER, RTA is liberated from RTB and retrotranslocated into the cytoplasm *via* the Sec61 complex (11). RTA is an RNA N-glycosidase that catalyzes the hydrolysis of a conserved adenine residue within the sarcin/ricin loop of 28S rRNA, resulting in the inhibition of protein synthesis (12, 13) and the activation of apoptosis (14). Alveolar macrophages are particularly sensitive to the cytotoxic effects and secrete an array of pro-inflammatory cytokines before undergoing apoptosis (8, 9, 15).

In this report we investigated the potential of a humanized monoclonal antibody (MAb) huPB10 to serve as a therapeutic in an established Rhesus macaques model of RT inhalation (3, 16). PB10 was first described as a murine MAb with potent-toxin neutralizing activity *in vitro* and *in vivo* (17). PB10 recognizes an immunodominant epitope situated at the apex of RTA, relative to RTB (18, 19). Chimeric (cPB10) and fully humanized (huPB10) versions of PB10 retain *in vitro* toxin-neutralizing activity and have been shown to passively protect mice against lethal dose of RT administered by injection or inhalation (20, 21). It was also demonstrated that huPB10 can rescue mice from intoxication if administered 4-6 h after exposure (21). Based on these preliminary findings, assessed the therapeutic potential of huPB10 in a well-established nonhuman primate (NHP) model of ricin aerosol challenge (16). The NHP is believed to be the model most representative of aerosolized exposure.

## Results

A total of 12 rhesus macaques (∼7 kg; range 3.8-10.2 kg) bred at Tulane National Primate Research Center (TNPRC) were randomly assigned to three experimental groups and then challenged with RT by small particle aerosol at a target dose of 18 µg/kg (**Table 1; Figure 1**). Animal studies were conducted in strict compliance with protocols approved by TNPRC’s Institutional Animal Care and Use Committee (IACUC). Group 1 (n=2) received intravenous administration of saline 4 h post challenge. Animals in groups 2 and 3 received huPB10 at 4 h or 12 h post challenge, respectively (**Table 1**). huPB10 was administered intravenously at a final dose of 10 mg/kg (**Table S1**). The macaques were subjected to whole body plethysmography and radiotelemetry over the course of the study. Animals surviving on day 14 post challenge were euthanized and subjected to complete necropsy and histopathological analysis. Serum and bronchial alveolar lavage (BAL) fluids were collected from the animals before and 24 h after exposure to RT.

**Table 1.** Experimental groups and outcome of ricin challenge

| Group | ID | kg | M/F<sup>a</sup> | RT<sup>b</sup> | huPB10 (µg/ml)<sup>c</sup> |  | TTD<sup>d</sup> | #/total |
| --- | --- | --- | --- | --- | --- | --- | --- | --- |
|  |  |  |  |  | serum | BAL |  |  |
| Control | DH27 | 9.7 | M | 14 | - | - | 28 h | 0/2 |
|  | JP56 | 8.4 | M | 26 | - | - | 30 h |  |
| +4 h huPB10 | LJ80 | 4.4 | M | 31 | 157 | 2.8 | + | 5/5 |
|  | LB98 | 4.9 | M | 29 | 177 | 2.7 | + |  |
|  | EM61 | 9.5 | M | 31 | 207 | 2.5 | + |  |
|  | IC70 | 5.9 | F | 26 | 225 | 3.6 | + |  |
|  | IL40 | 6.1 | F | 33 | 147 | 3.2 | + |  |
| +12 h huPB10 | JM39 | 9.4 | M | 22 | 205 | 42.8 | 72 h | 1/5 |
|  | DR61 | 10.2 | F | 24 | 265 | 3.9 | + |  |
|  | KP48 | 5.8 | M | 30 | 203 | 25.7 | 48 h |  |
|  | KR49 | 5.7 | M | 29 | 151 | 3.1 | 49 h |  |
|  | LB26 | 3.8 | M | 29 | 191 | 11.3 | 44 h |  |

<sup>a</sup>, sex of animal; <sup>b</sup>, ricin toxin (RT) dose (µg/kg) received per animal; <sup>c</sup>, huPB10 in serum and BAL from samples collected at 24 h post challenge. The average in serum was 182.6 µg/ml and 203 µg/ml for the +4 and +12 h groups, respectively. In the BAL, the was 3.04 µg/ml and 17.3 µg/ml, respectively. Times of huPB10 administration are approximate (~ 30 minutes of point estimate); the logistics of performing slow intravenous infusion of huPB10 in monkeys initiated and was times at 4 hours from each of the individual ricin aerosol challenge events and was performed in sequence; <sup>d</sup>, time to death (TTD); +, indicates survival for 21 day duration of study.

**Figure 1.**
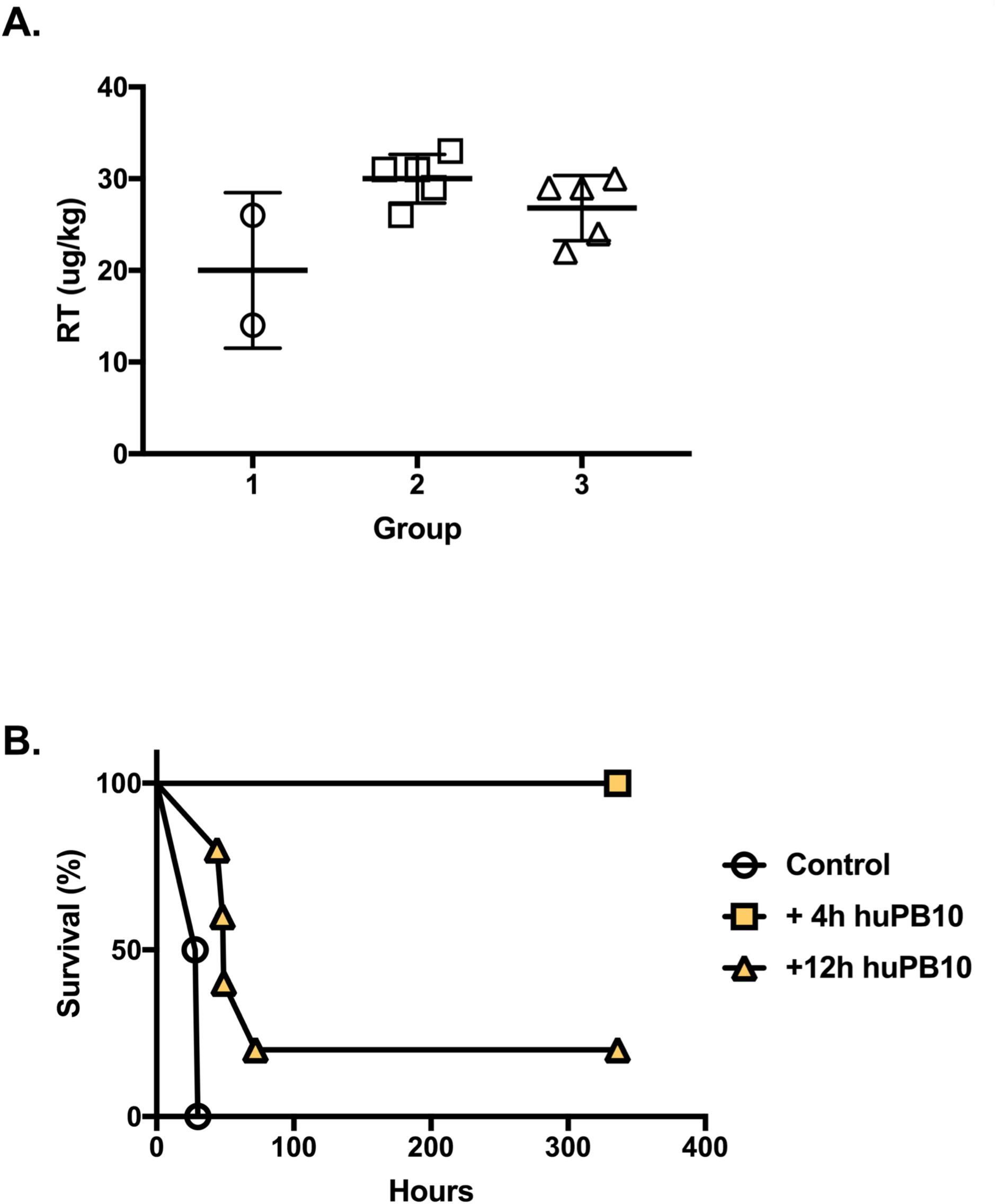
Effect of huPB10 on the survival of animals exposed to aerosolized RT. A. Animals were exposed to aerosolized RT aerosol and the individual RT doses, expressed in μg/kg inhaled, and clustered by treatment group, show minimal variation in dosing, bars represent group mean ± standard deviation; the aerosol target dose of 18 µg/kg represented by segmented line. B. Macaque survival is represented by survival curve. The two control animals succumbed to intoxication by day 2; treatment with huPB10 at 4 h post ricin challenge resulted in 5/5 survival, while treatment with huPB10 at 12 h resulted in only 1/5 survivors.

Control animals (n=2) succumbed to RT toxicosis within 36 hours of exposure. (**Table 1; Figure 1**). The clinical progression of RT-induced intoxication of the control animals was the same as in previous studies (3). Approximately 12-16 hours after exposure, animals displayed reduced activity and fever (**Figure 2**). Clinical examination conducted 24 h post exposure revealed respiratory complications, including bilateral congestion and crackles, with dyspnea and tachypnea. Arterial oxygen ranged from 75-85%. The clinical state of the animals continued to decline over the subsequent several hours, manifested by a marked drop in normal activity and cyanotic mucous membrane.

**Figure 2.**
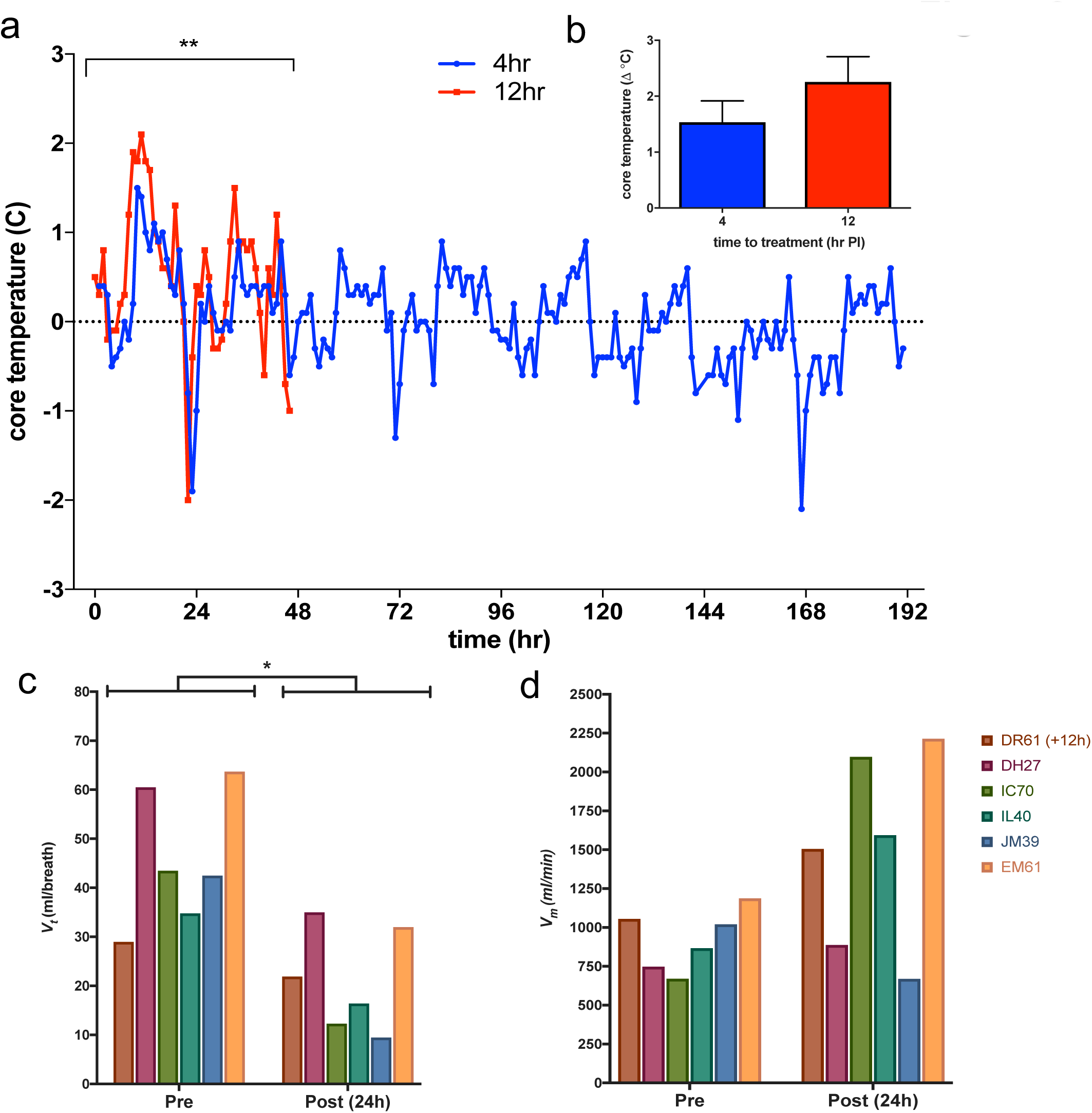
Physiological response to RT. A. Radiotelemetry of core temperature of rhesus macaques treated with huPB10 either at 4 h or 12 h post-exposure to aerosolized RT. Continuous monitoring showed significant differences (*P*<0.005) between the initiation and tempo of pyrexia that was dependent on the time to treat. B. Fever intensity showed differences between the relative change (increase) based upon time to treat. Respiratory function measured by whole-body plethysmography. Respiratory function was measured *via* head-out conductance plethysmography prior to and at timed intervals post challenge. C. Significant changes (*P*<0.05) in post-exposure group tidal volume was observed among animals rescued from ricin intoxication, (D) although compensatory changes in frequency resulted in minimal observable changes in minute volume, with some animals producing increased minute volumes +24 h post-exposure to ricin.

In contrast, all five of the Rhesus macaques that received huPB10 at 4 h post-ricin challenge time point survived (*P<0.01* compared to controls) and remained otherwise normal for the duration of the three-week post-exposure observation period (**Table 1: Figure 1**). Compared to the control animals, the five macaques in Group 2 presented with only minimal signs of distress: mild dyspnea, minimal tachypnea and increased lung sounds upon physical examination at 24 h post-exposure. The animals displayed no such symptoms upon physical examination on day 7.

Finally, only one of the five animals in Group 3 that received huPB10 at 12 h survived RT challenge (**Table 1; Figure 1;** *P<0.01* and *P<0.05* compared to controls and +4 h group, respectively). The remaining four animals succumbed to RT intoxication between 44 and 72 h post challenge and followed a clinical course similar to the control animals. HuPB10 was detected in the sera and BAL fluids collected at 24 h post RT challenge, indicating that biodistribution of huPB10 was similar between groups 2 and 3.

Gross examination of the lungs from sham-treated control animals showed coalescing hemorrhage with frothy exudate marked by fibrin in lung parenchyma (**Figure S1**). The lungs of control animals were grossly described as fibrinosuppurative bronchointerstitial pneumonia with pulmonary edema and bronchial epithelial necrosis, with severe fibrinosuppurative lymphadenitis in the bronchial lymph nodes. The wet weights of control animal lungs were >150 grams in contrast to a normal wet lung weight of ∼30-40 grams in naïve animals of approximately the same body weight. Histologically, the lungs of sham-treated animals showed hallmark inflammation consistent with RT-induced injury, characterized by marked edema, corresponding hemorrhage, and numerous infiltrates. The pathological outcome of the four animals in Animals in Group 3 that succumbed to RT intoxication resembled the control animals. There was extensive pulmonary congestion, edema, inflammation with infiltrates, and punctate hemorrhage evident. Gross lung weights were similar to RT-challenged control animals (>140 g) with clear signs of hemorrhage.

Pathological analysis of the lung tissues collected at the time of euthanasia (day 21 post challenge) from the five survivors in Group 2 that had been treated with huPB10 at 4 h and the single animal in Group 3 (DR61) revealed evidence of chronic inflammation and a distinctive fibrosis proximal to the respiratory bronicholes, reminiscent of past studies in which animals had received sub-lethal exposures to RT (22).

Sera and BAL fluids collected before (day −7) and 24 h post RT challenge were subjected to analysis with a 29-plex cytokine/chemokine/growth factor Luminex array as a means to assess the impact of huPB10 on local and systemic inflammatory responses. In the sera of the two control animals, there were 12 cytokines/chemokines that were significantly different in serum post-versus pre-challenge; 11 cytokines were increased, while one (IL-8) was decreased (**Figure 3A; S2**). Most notable were IL-6 (∼500-fold increase) and IL-1RA (∼256-fold increase). VEGF was also elevated, possibly reflecting response to pulmonary insult. Analysis of sera from animals that received huPB10 at 4 h post challenge indicated that only four cytokine/chemokines were significantly different from pre-challenge levels (3 up;1 down), including a ∼4-fold increase in IL-6. IL-1RA and VEGF levels were unchanged. In animals that received huPB10 at 12 h after RT challenge, a total of six cytokines/chemokines changed relative to pre-challenge levels, although the magnitude of these changes was lower than that if the control animals, possibly reflecting a dampening of the inflammatory response as a consequence of huPB10 intervention. Principle component analysis (PCA) of fold-change in cytokine/chemokine levels from serum samples from all 12 monkeys did not reveal any clustering by experimental group or survival (**Figure S3A-C**), signifying that serum inflammatory responses are not indicative of experimental outcome.

**Figure 3.**
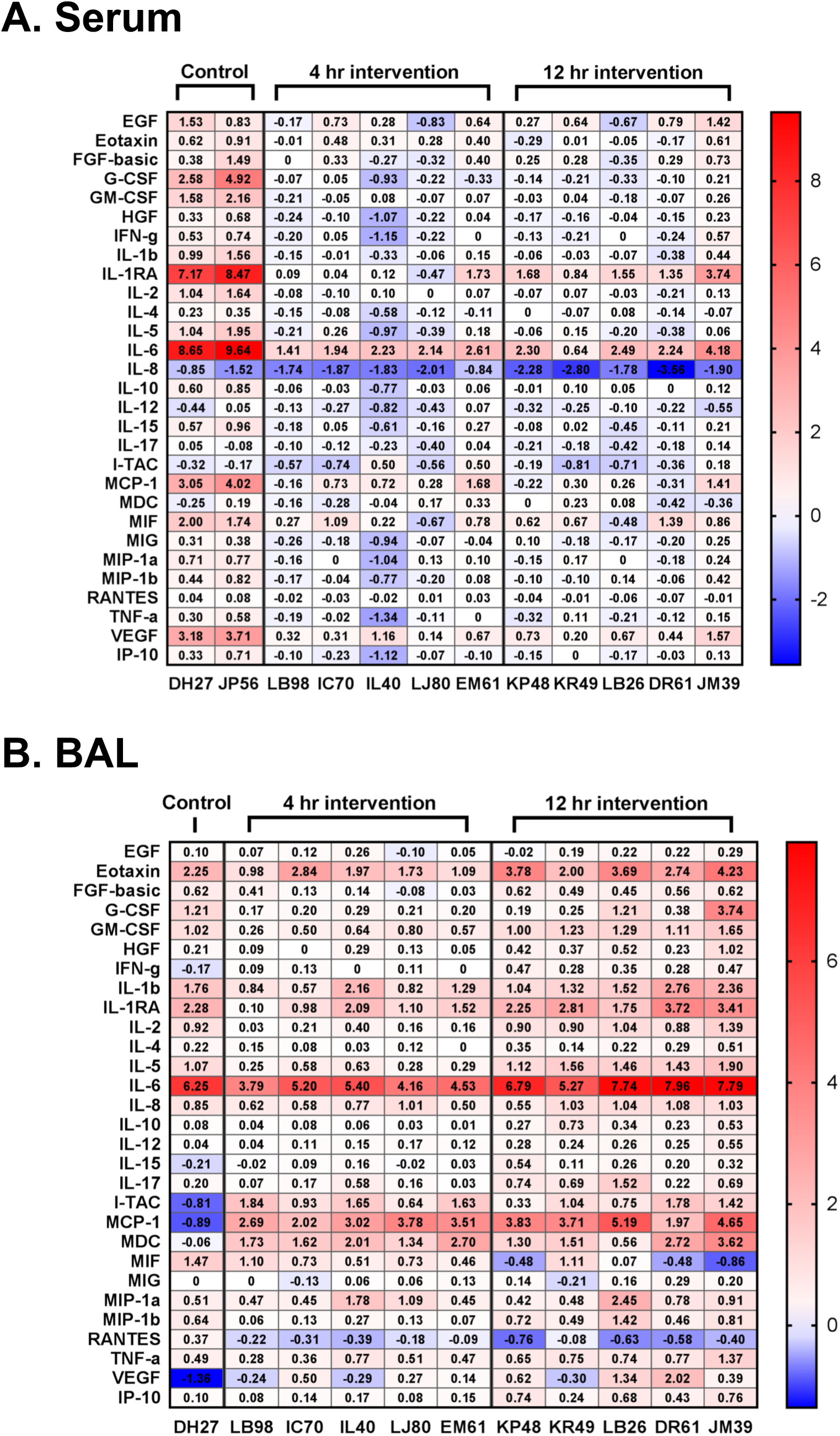
Cytokine profiles of serum and BAL in RT challenged Rhesus macaques. Heatmaps present log_2_-fold change of cytokines in (A) sera and (B) BALs, comparing samples collected 24 hours post-challenge to those collected 7 days pre-challenge. Animals are separated into groups for controls, 4-hour, and 12-hour huPB10 interventions. Increases in cytokine levels are shown in red, while decreases are shown in blue. Specific values for the fold changes are shown within each cell and based on single 29-plex analysis. A subset of cytokine changes were confirmed using BD™ CBA Human Inflammatory Cytokines Kit.

The impact of huPB10 intervention was much more apparent in BAL fluids than in sera (**Figure 3B**). Within group 2, a total of 21 cytokines/chemokines were significantly elevated following RT challenge (as compared to pre-challenge levels), with IL-6 being the most pronounced. In group 3, there were 25 cytokines/chemokines that were significantly elevated compared to pre-challenge levels. Seven cytokines/chemokines differed between the groups 2 and 3 with the most notable being IL-6, which was 32-fold elevated in group 2 and 181-fold elevated in group 3. PCA successfully clustered animals by both group and survival status, demonstrating that the localized cytokine responses in the BAL are more closely related to survival than the systemic responses (**Figure S3D-E**). The relative contributions of the different cytokines/chemokines responsible for segregating the animals into clusters are shown in **Figure S3F**.

## Discussion

The results of this study constitute a significant advance in longstanding efforts to develop effective medical countermeasures against RT inhalation (2, 23). Foremost, it is the first demonstration in NHPs that a MAb can rescue animals from the lethal effects of aerosolized ricin toxin exposure. Studies in primates are important because anti-RT products for humans must adhere to the Food and Drug Administration’s (FDA) Animal Rule; human challenge studies with RT are obviously unethical. The well-established model of aerosolized RT challenge in Rhesus macaques was a prerequisite for the therapeutic study conducted herein (3, 16, 24). Comparative models in mice (6, 8, 25) and swine (10) may also be important for therapeutic development under Animal Rule guidelines. The fact that all five animals in group 2 (+4 h huPB10 intervention), and one animal in group 3 (+12 h huPB10 intervention), survived exposure to RT indicates that a significant proportion of RT remains accessible to huPB0 in the alveolar space and/or interstitial fluids for hours after inhalational exposure. This finding is consistent with reports in mice with huPB10 and other anti-RT MAbs (6, 7, 21, 26), but surprising all the same considering the extraordinary sensitivity of the lung mucosa to the effects of toxin exposure (27).

We postulate that high dose intravenous delivery of huPB10 results in the accumulation of huPB10 within the lung mucosa and alveolar space where it can engage free (soluble) or receptor-bound RT. Indeed, IV administration of recombinant anti-viral IgG1 MAbs results a corresponding linear distribution of antibodies lung (28, 29). Whether transudation or active transport of serum antibodies into the lung mucosa is triggered by RT exposure has not been evaluated. Nonetheless, once in the lung mucosa, huPB10 presumably limits toxin uptake into target cells (e.g., macrophages and airway epithelial cells) and/or interferes with ricin intracellular transport in the event that endocytosis of RT-antibody complexes should occur (30, 31). Work by Magun and colleagues made it clear more than a decade ago that protecting alveolar macrophages is paramount in limiting toxin-induced lung damage (9, 32).

It is worth underscoring the value of the 29-plex monkey cytokine/chemokine/growth factor array in not only identifying local inflammatory markers like IL-6 that arise following RT exposure but also enabling through PCA the unbiased clustering of specific animals based on survival or experimental group. Identifying an inflammatory “fingerprint” associated with RT exposure has obvious applications in diagnostics, especially in the context of biodefense where early clinical symptoms following exposure to different toxins and pathogens may in fact be indistinguishable (33). As noted above, IL-6 levels were particularly elevated following RT exposure, which is consistent with what has been observed in mice (34). IL-6 is a particularly potent driver of pulmonary inflammation and could very well be a major contributor to RT-induced pathology in conjunction with RT’s other properties, including agglutinin activity and the capacity to induce vascular leak syndrome (5, 35). We recently reported that human lung epithelial cell lines preferentially secrete IL-6 following RT exposure, especially in the presence of pro-apoptotic factors like TRAIL (36). It has been suggested that anti-inflammatory agents may extend the therapeutic window of anti-RT MAbs by suppressing bystander tissue damage (26). Whether such directed immunotherapies would have utility in the context of potent toxin-neutralizing antibody like huPb10 remains to be seen. At least early intervention with huPB10 was sufficient to render RT effectively inert within the context of the lung and it seems unlikely that supplementing treatment with anti-inflammatory agents would afford much additional benefit.

## Methods

### Ricin toxin and huPB10

Purified ricin toxin derived from castor beans (*Ricinus communis*) was produced as previously described (37). HuPB10 was expressed using a *Nicotiana benthamiana*-based manufacturing platform. The properties of huPB10 used in this study are shown in **Table S1.**

### Animal care and use

Rhesus macaques were born and housed at the Tulane National Primate Research Center (Covington, LA), which is US Department of Agriculture-licensed and fully accredited by the Association for Assessment and Accreditation of Laboratory Animal Care (AAALAC). Aerosolization, dosing and delivery of RT were performed as described (16). The LD_50_ of ricin is 5.8 µg/kg body weight and the target dose for this experiment was set at the equivalent of three LD_50_s (≈18 µg/kg). The mean inhaled dose of ricin across all animals was 4.4 ± 1.4 LD_50_s. At 4 h or 12 hours post-exposure, designated animal groups received a single intravenous administration of huPB10 by slow infusion at an individualized unit dose of 10 mg/kg. Sham-treated animals received saline at the 4h time point. Treated animals were observed for signs of adverse reactions to the MAb during administration and throughout the anesthesia recovery period. Animals were bled just before and 24 h following aerosol challenge. Blood was also collected when the animals either succumbed to intoxication or 21 d after challenge, when the experiment was terminated. Animals determined to be in respiratory distress and those that survived for 21 d after exposure to ricin were euthanized by an overdose of sodium pentobarbital, consistent with the recommendation of the American Veterinary Medical Association’s Panel on Euthanasia, and submitted for necropsy. All methods were approved by the Tulane University’s IACUC. After gross necropsy, tissues were collected in neutral buffered zinc-formalin solution (Z-Fix Concentrate, Anatach). Tissues were processed, sectioned, and stained as previously described (3).

### Statistics

Statistical analysis was carried out with GraphPad Prism 6 (GraphPad Software, La Jolla California USA). The difference in outcomes between groups was determined by Fisher’s exact test (two-tailed) and the mean survival times after exposure to RT were compared by log-rank analysis of Kaplan–Meier survival curves. The statistical significance of the effects of ricin challenge and huPB10 intervention on cytokine levels were analyzed with two-way repeated measures ANOVAs in both serum and BAL, with the repeated measures being pre- and post-exposure status, and treatment group as the independent measure. Resulting p-values were corrected with the Benjamini, Krieger, and Yekutieli method to control false discovery rate. All analyses were performed on log^2^ transformed raw fluorescent intensity values to avoid the need to censor values. PCA analysis was performed using singular value decomposition with the R package FactoMineR (38). Heatmap construction using raw fold change values was done using GraphPad Prism 6. Hierarchical clustering and scaled heatmap construction was completed with the R package pheatmap (39).

## Author contributions

NB, OB, DK, MP, KW and ESV generated reagents; PJD, LDM, YR, and CJR conducted animal studies and animal tissue/sample analysis; DJE performed statistical analysis; CJR, NJM, and LZ are responsible for experimental design and CJR, NM and ESV prepared the manuscript.

## Conflicts of interest

NB, OB, DK, MP are Mapp Biopharmaceutical employees and shareholders. KW and LZ are employees, shareholders, and co-owners of Mapp Biopharmaceutical. The remaining authors declare no conflicts of interest.

## Acknowledgements

This work was supported by R01AI098774 to LZ and Contract No. HHSN272201400021C to NJM from the National Institutes of Allergy and Infectious Diseases, National Institutes of Health (NIH), as well as the National Center for Research Resources and the Office of Research Infrastructure Programs (ORIP) of the NIH through grant OD011104 at the Tulane National Primate Research Center to CJR. The content is solely the responsibility of the authors and does not necessarily represent the official views of the National Institutes of Health. The funders had no role in study design, data collection and analysis, decision to publish, or preparation of the manuscript.

## Supplemental Material

### Supplemental Methods

#### Animal Husbandry and Telemetry

Rhesus macaques were born and housed at the Tulane National Primate Research Center (Covington, LA), which is US Department of Agriculture-licensed and fully accredited by the Association for Assessment and Accreditation of Laboratory Animal Care. Subcutaneous radiotelemetry transmitters combined with sensors capable of detecting biopotential signals of an electrocardiogram as well as thermistor-type sensors capable of detecting temperature signals (T34G-8; Konigsberg Instruments) were surgically implanted under aseptic conditions in 8 of the 12 vaccinated macaques and both control macaques before the start of the study. Animals determined to be in respiratory distress and those that survived for 21 d after exposure to ricin were euthanized by an overdose of sodium pentobarbital, consistent with the recommendation of the American Veterinary Medical Association’s Panel on Euthanasia, and submitted for necropsy. All methods were approved by the Tulane Institutional Animal Care and Use Committee (IACUC).

#### Treatment of Rhesus macaques

At 4 h or 12 hours postexposure, designated animal groups received a single intravenous administration of huPB10 by slow infusion at an individualized unit dose of 10 mg/kg. Sham-treated animals were administered saline for injection at the four-hour time point. Treated animals were observed for signs of adverse reactions to the antibody during administration and throughout anesthesia recovery period. Animals were bled just before and 24 h following aerosol challenge. Blood was also collected when the animals either succumbed to intoxication or 21 d after challenge, when the experiment was terminated.

#### RT aerosolization, dosing, and calculation

Aerosolization, dosing and delivery of ricin were performed as described (16). Inductive plethysmography that measures volume of air breathed by each individual animal per minute was performed just before the ricin exposure. Ricin was dissolved in 10 mL sterile phosphate buffer saline to the desired concentration for each animal based on plethysmography data obtained 2 d before the exposure. Aerosols were generated directly into a head-only chamber using a Collision three jet-nebulizer (BGI) with fully automated management control system (Biaera Technologies, Hagerstown, MD) all within a Class III biological safety cabinet housed within the TNPRC high-containment (BSL-3) laboratories. The nebulizer operated at 18 lb/inch^2^ equating to a flow of 7.5 L/min and produced 3.0E+04 particles per cc with a mass median aerodynamic diameter of ∼1.4 µm. Each discrete aerosol exposure lasted 10 minutes (per animal). Air samples were continuously obtained during the exposure and the protein concentrations of these samples were determined using a micro-BSA protein assay kit (Thermo Scientific). The aerosol concentrations were determined and the inhaled dose of RT for each animal was calculated by multiplying the empirically determined aerosol exposure concentration (µg/liter of air) in the chamber by volume of air estimated to have been breathed by the animal (via results of plethysmography just before exposure). The LD_50_ of ricin is 5.8 µg/kg body weight and the target dose for this experiment was set at the equivalent of three LD_50_s (≈18 µg/kg). The mean inhaled dose of ricin across all animals was 4.4 ± 1.4 LD_50_s.

#### Statistics

Statistical analysis was carried out with GraphPad Prism 6 (GraphPad Software, La Jolla California USA). The difference in outcomes between groups was determined by Fisher’s exact test (two-tailed) and the mean survival times after exposure to ricin were compared by log-rank analysis of Kaplan–Meier survival curves. The statistical significance of the effects of ricin challenge and huPB10 intervention on cytokine levels were analyzed with two-way repeated measures ANOVAs in both serum and BAL, with the repeated measures being pre- and post-exposure status, and treatment group as the independent measure. Resulting p-values were corrected with the Benjamini, Krieger, and Yekutieli method to control false discovery rate. All analyses were performed on log^2^ transformed raw fluorescent intensity values to avoid the need to censor values. PCA analysis was performed using singular value decomposition with the R package FactoMineR (38). Heatmap construction using raw fold change values was done using GraphPad Prism 6. Hierarchical clustering and scaled heatmap construction was completed with the R package pheatmap (39).

#### Tissue Collection, Histological Analysis, and Special Stains

After gross necropsy, tissues were collected in neutral buffered zinc-formalin solution (Z-Fix Concentrate, Anatach). Tissues were processed, sectioned, and stained as previously described (3).

## Supplemental Table

**Table S1.**
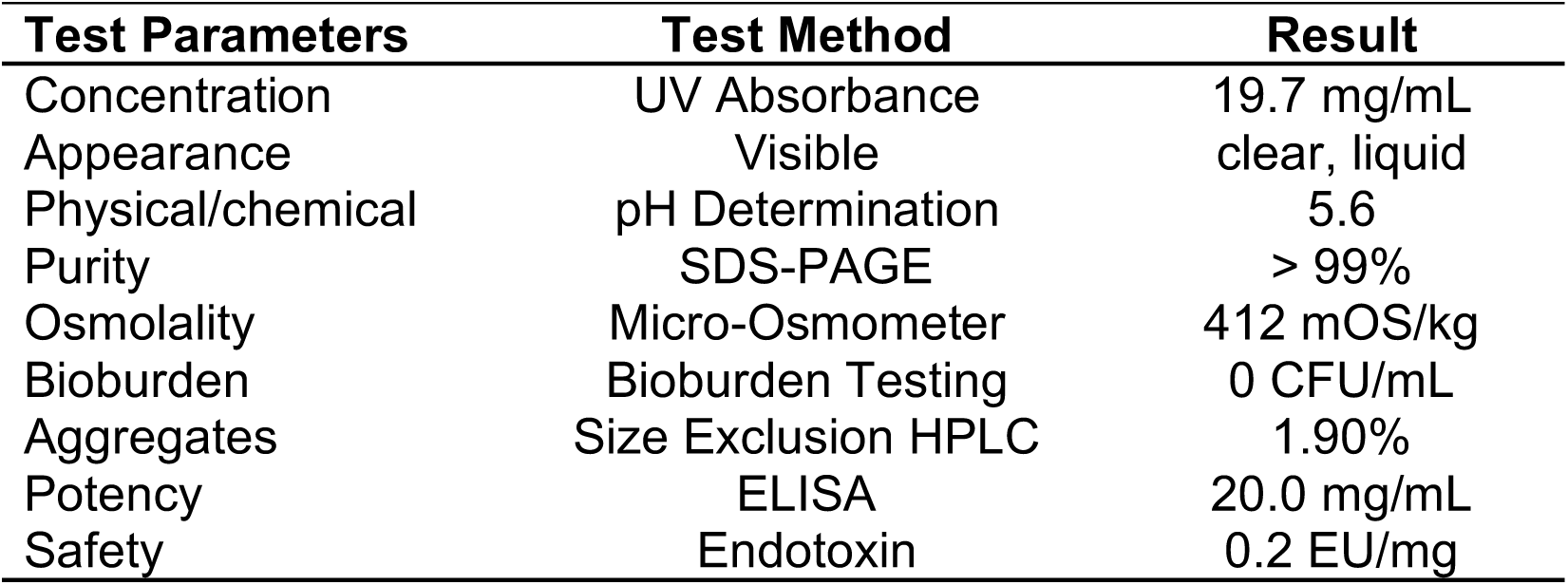
Characteristics of huPB10 used in study

**Figure S1.**
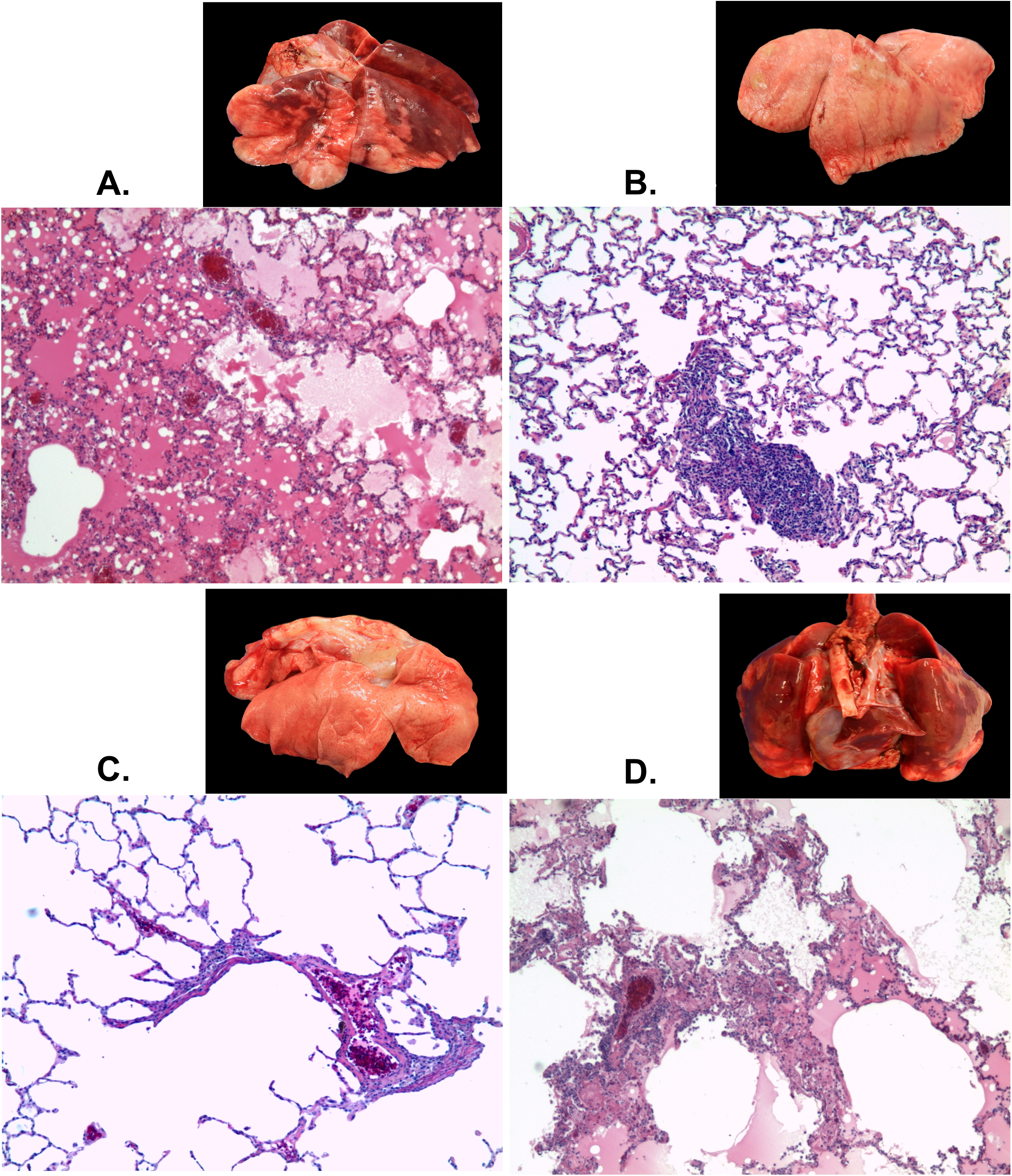
Gross pathology associated with RT exposure and huPB10 intervention. The lungs of sham-treated animals (Figure 3a) show hallmark inflammation, marked edema, corresponding hemorrhage, and numerous infiltrates; grossly, wet weight at 3x normal with coalescing hemorrhage. Animals treated with huPB10 at 4 h (Figure 3b & 3c) show remarkably little pulmonary damage, with occasional infiltrates; lung wet weights were essentially normal. Animals treated with huPB10 at 12 hours (Figure 3d) demonstrated mild to moderate inflammation with infiltrates, edema, and punctate hemorrhage evident; gross lung weight were approximately 2x with clear signs of hemorrhage. Original magnification at 10x

**Fig. S2.**
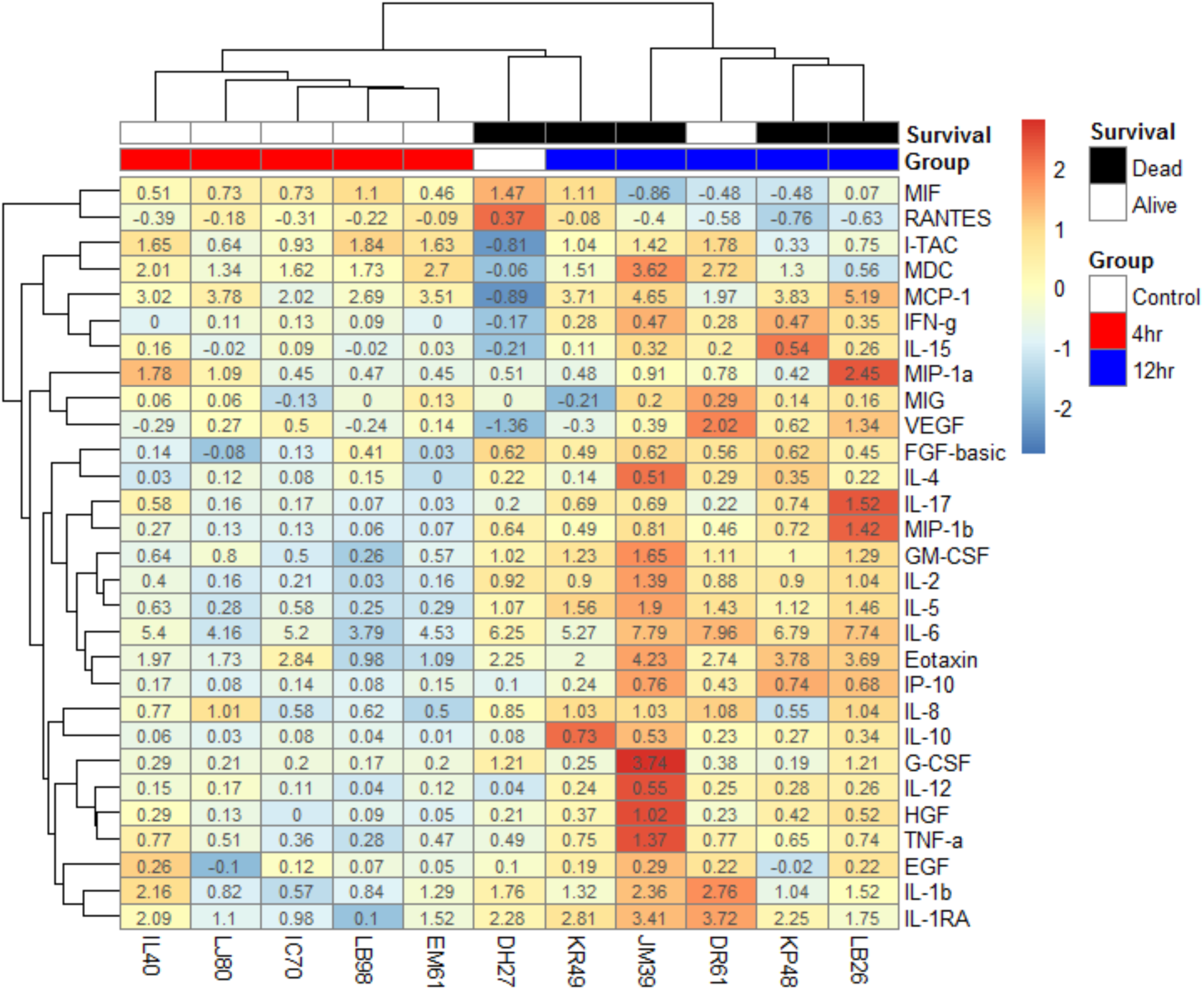
Scaled heatmap and cluster analysis of log2-fold changes in cytokine levels in individual macaques following ricin challenge. Heatmaps visualizing log2-fold changes in cytokines in serum (A) and BAL (B). Both individual animals and cytokines are arranged by hierarchical clustering, represented by dendrograms on top and to the right of the heatmaps, respectively. Surviving animals are marked in white in the top bar above the heatmap, while dead animals are in black. In the bar below this, the 4-hour group of animals are marked in red, the 12-hour group in blue, and the control group in green. Fold change values are centered and scaled for each cytokine by first subtracting the mean fold change from each value and then dividing by the standard deviation of that cytokine.

**Fig. S3.**
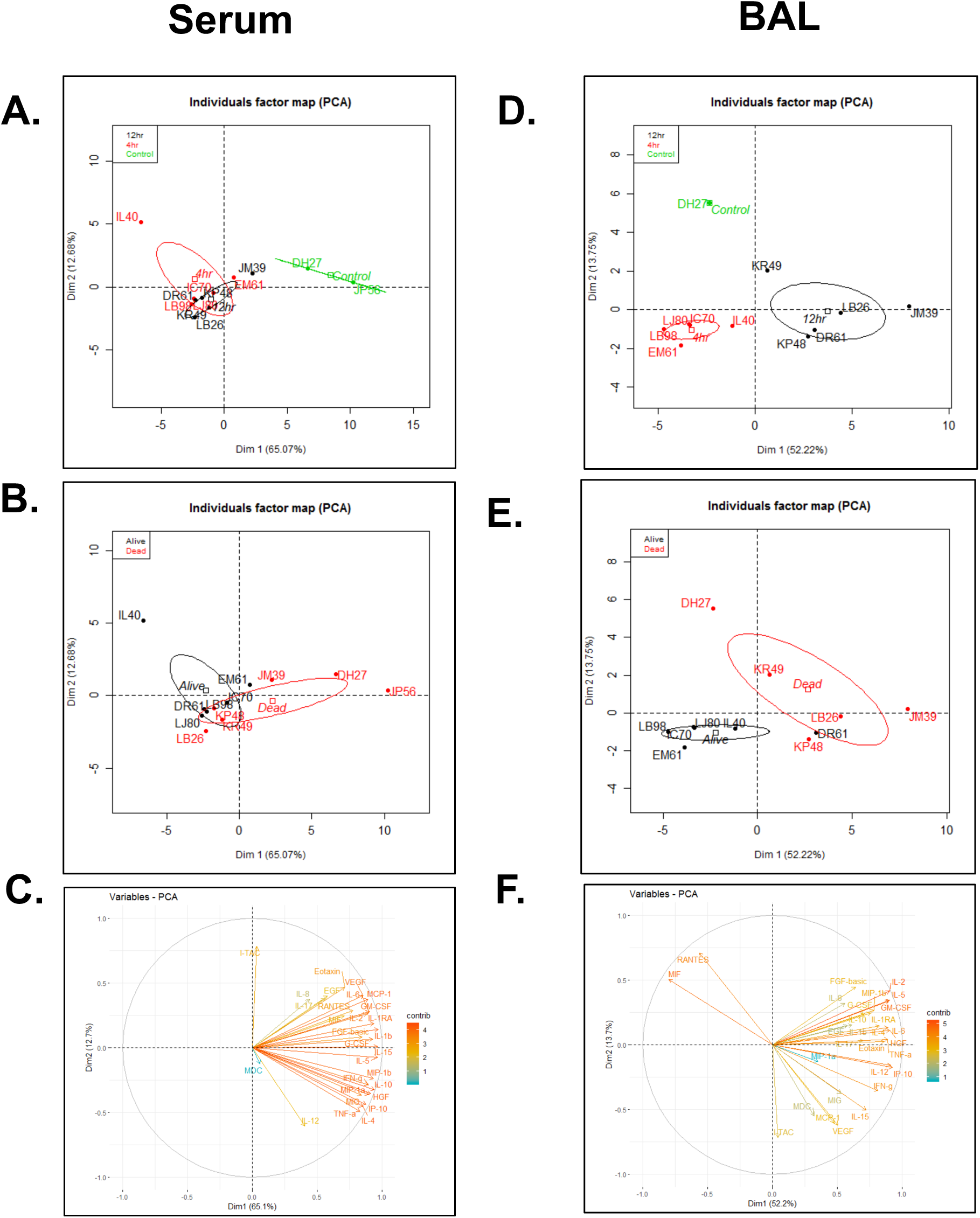
Principal component analysis of cytokine fold changes following RT challenge. Scatter plots of first 2 principal components of log2-transformed cytokine fold changes in A-C) BAL and (D-F) serum from all animals included in the study, calculated used singular value decomposition. Each dot represents an individual monkey; red for those in the 4-hour group, black for the 12-hour group, and green for the control group. 95% confidence ellipses of the group means are represented in the color of the group they correspond to. Eigenvectors for (C) BAL and (F) and analyses are colored to show the percent contribution of each variable to the principal components.

